# A preliminary study of the calcification reason of pit mud in Chinese Strong-flavour liquor

**DOI:** 10.1101/319996

**Authors:** Bin Chen, Deliang Wang, Yinzhuo Yan

## Abstract

Pit mud play a key role in the brewing process of Chinese Strong-flavour liquor. In this study, the organic acids and metal ions was determinated by Ion Chromatograph and Inductively Coupled Plasma Mass Spectrometry, respectively. The diversities and structures in pit mud were analyzed by using high-throughput sequencing based on Illumina Miseq platform. The results showed that the lactic acid in calcified pit mud was about 10 times higher than quality pit mud, and the calcium ions was about 20 times higher than quality pit mud. It could found a strange phenomenon, the calcified pit mud had more pyruvic acid and quality pit mud basically does not had this kind of material. The data analysis shown that the pit mud were detected prokaryotes 908 strains and the eukaryotes 226 strains. It was found that the Lactobacillus and Prevotella in calcified pit mud (the total of both increase 11 times) had a higher percentage. It can be guess that the lactic acid and metal ions would form lactic acid salt compounds and those compounds could affect the growth and reproduction of microbes in pit mud. This study provided a good comparison of physical and chemical properties between the calcified pit mud and the quality pit mud, which was significant for the maintenance of pit mud and the quality of liquor. Because of the first study for pit mud by using high-throughput sequencing, it could provide theoretical basis for improving microbes in pit mud.

## Importance

This article introduced the study of pit mud, it found that there were many difference between the calcified pit mud and quality pit mud.

This study provided a good comparison of microbial diversities and structures between the calcified pit mud and the quality pit mud, which was significant for the maintenance of pit mud and improved the quality of liquor.

To the best of our knowledge, this is the first report about the microbial community structure and diversity in calcified pit mud and quality pit mud by using high-throughput sequencing, which will help us to improve the quality of pit mud, solve the problem of calcification and enhance the quality of the Strong-flavor liquor.

## Introduction

Chinese liquor is one of the six most famous distilled alcoholic beverages in the world and is very popular in China. In general, it is mainly classified into the six types: Strong-flavour, Light-flavour, Sauce-flavour, Sesame-flavour, Rice-flavor and other different flavours. The most popular type is Strong -flavour liquor in the market, which was about 70% of all types *(1).* It uses pit mud as the fermentation cornerstone, which is a fermenting agent and has a significant impact on the flavour of the product. Therefore, high quality aged pit mud has the reputation of ‘National treasure mud’ and ‘Microbial gold’ in China. However, with the increase of pits mud aging, the main question is the calcification, which will leads to the pits mud become more and more rigid and even appear the white crystalline. The crystalline will directly affects the microbial diversities and structures of pit mud. It is a valuable industry resource, but the loss and lack of this resource are also obvious recently. So it is necessary to understand the reason of calcification, especially the imparity of the two types. The study would focus on the research of microbial structure.

Analyzing the structures and diversities of the microbial community has always been a big challenge for Chinese microbiologists. Pit mud quality, according to its sensory characteristics, can be divided into three grades: degraded, normal, and high quality. However, the relationship between pit mud microbial community and pit mud quality is poorly understood *(2).* In the latest thirty years, most Chinese microbiologists used culture-dependent and culture-independent methods to analyze the dominating microbes in pit mud. In 1991, a strain of *Clostridium prazmowski* was found as the main microorganism in *Wuliangye* pit mud by the culture-dependent method *(3).* With the progress in technology, there are many approaches to study the microorganism in the pit mud, such as denaturing gradient gel electrophoresis (DGGE), phospholipids fatty acid (PLFA), High-throughput sequencing. Recently, Clostridiales, Lacotobacillales and Bacillales were reported as the main bacterial microorganisms and *Pichia anomala* was the main yeast in pit mud by combined DGGE and PLFA analyses *(4).* Lately, Liang H et al *(5),* found that there were some significant differences in the microbial community structure between aged and aging pit mud by the methods of PCR-DGGE and qPCR. Li C et al *(6),* found clostridium butyricum is an important fragrance-producing bacterium in the traditional Chinese flavor liquor-making industry. But the technology of high-throughput sequencing was not used currently. It was a cost-expensive method for knowing the characterization of pit mud microbial communities, but these analysis methods were very effective. The analysis can get millions of DNA sequence and transcriptome sequencing or genome sequencing. So it is cost-effective for the study of microbial diversities and structures. In this study, Illumina MiSeq platform was used to evaluate the differences in calcified pit mud and quality pit mud. To the best of our knowledge, this is the first report about the microbial community structure and diversity in calcified pit mud and quality pit mud by using high-throughput sequencing.

Through the study of the physical and chemical properties and microbial structure of the two kinds of pit mud, it was found some difference. Hence, the structure and composition of microbes studied by modern science and technology has become a new subject in liquor-making industry, which will help us to understand the pit mud and even solve the problem of calcification or enhance the quality of the Strong-flavour liquor.

## Materials and Methods

### Sampling

In this study, the quality and calcified pit mud samples were collected from *Yanghe* and *shuanggou* manufactory, which were located in Jiangsu province in China. S5JD1, S5JB1, S5JD3 and S5JD4Ca were collected from the *Shuanggou.* They were all quality pit muds, except the last sample. YH1JN, YH11JD, YH11BCad and YH11BCau were collected from the *Yanghe.* The first two were the quality pit muds and the last two were the calcified pit muds. Approximately 20 g of each sample were transferred to sterile polyethylene bags and stored at -20 °C until analyzed.

### The determination of organic acids in pit mud

For the determination of organic acids in pit mud, this experiment adopts ISC3000 ion chromatograph, concrete operation as follows, (1) Take the pit mud 5 g, dissolved in 25 mL ultrapure water, after fully oscillation, 2 000 rpm centrifuge for 10 min. (2) Through 0.45 water phase membrane to filter and diluted 100 times, then testing by ISC3000 ion chromatograph. Instrument set conditions as follows: pillar type IONPAC^®^ AG11-HC 4 mm×50 mm, pillar temperature 30°C, nitrogen pressure 6 kPa. In order to guarantee the authenticity of the sample data, it was necessary to every four samples to join a standard sample (the lactic acid concentration 50 μg/L).

### The determination of metal ions in pit mud

For the determination of metal ions in pit mud, this experiment adopts NexION 350 Inductively Coupled Plasma Mass Spectrometer (ICP-MS), concrete operation as follows, (1) Take some pit mud, air dry for one week and grinding, then through 200 mesh sieve. (2) Accurate weighing 0.2 g (Accurate to 0.001), then placed in digestion jar, add 4 mL of nitric acid (HNO_3_), 2 mL H_2_O_2_, 2 mL hydrofluoric acid (HF), finally put the digestion jar to microwave digestion instrument for 1 h. (3) After digestion, add 1 mL perchloric acid (HClO_4_), then placed on the heating plate, evaporation and take the remaining 1 mL. (4) Digestion jar samples transferred into 50 mL volumetric flask, then constant volume by 1% HNO_3_. (5) With 1% HNO_3_ solution diluted 100 times, then testing by ICP-MS.

### DNA extraction

Pit mud samples were washed three times with PBS solutions. The total DNA (prokaryotes and eukaryotes) from each sample was extracted by FastDNA SPIN Kit for Soil (MP Biomedicals, USA). The extraction process should according to the manufacturer’s instructions and the total DNA was stored at -80°C until further analysis. DNA concentration and purity were determined by a spectrophotometer (UV–VIS, SHIMADZU, Japan) and finally diluted to 50 ng/μL *(7).*

### PCR amplification

V4 regions of the 16S rDNA gene were amplified using the universal primers 515F and 806R *(8).* The fragments of ITS1 region from eukaryotes were amplified using the universal primers ITS1F and ITS1R *(9).* The PCR system were according the methods of Gerard *(10).* The conditions of the prokaryote amplification were according the methods of Luo *(1*). But for the eukaryotes, as follows, initial denaturation of DNA at 95 °C for 5 min, and 45 cycles of denaturation at 95°C for 30 s, then annealing at 50°C for 30 s and extension at 72°C for 1 min; at last, a final extension at 72°C for 5 min. Following successful amplification (identified by agarose gel electrophoresis), amplicon libraries were sequenced using Illumina MiSeq platform by Novogene Bioinformatics Technology Co. Ltd (Beijing, China).

### Data analysis

Due to the short read lengths and the overlap DNA fragments, FLASH (Fast Length Adjustment of SHort reads) which was a very fast software tool to find the correct overlap between paired-end reads and extend the reads by stitching them together was applied to solve those problem *(11).* The concrete was merge pairs of reads when the original DNA fragments were shorter than twice the length of reads. Then according to the unique barcode of each sample, the sequencing reads were assigned to each sample *(12).*

For the part of taxonomy, sequences were analyzed with the QIIME software package <u>(http://qiime.org/)</u> which supports a wide range of microbial community analyses *(13)* and UPARSE pipeline <u>(http://drive5.com/uparse/)</u> which picking operational taxonomic units (OTUs) through making OTU table *(14).* Sequences were assigned to OTUs at 97% similarity and also be compared with the Greengenes database *(15).* The representative sequences were picked for each OTU and used the RDP (Ribosomal Database Project) classifier to assign taxonomic data to each representative sequence *(16),* in addition to custom Perl scripts to analyze alpha diversity. The alpha diversity was used to determine whether cover all stains *(8*).

## Results

The pit mud plays an important role in the quality of Strong-flavour liquor. However, at present most manufactories find that the calcification of pit mud appears frequently, which directly affect the quality of liquor. This experiment mainly study physical and chemical properties and microbial structure of pit mud. Here were the results.

### The organic acids analysis of the two kinds pit mud

Because of the common acid of pit mud and it mainly from organic acid salts, especially the calcified pit mud, so it was necessary to find out what kind of organic acid to make it. Nine kinds of main organic acids were identified, the results were shown in table 1. From the table, it can be found that the highest organic acid was lactic acid in each samples. The lactic acid content of calcified samples (S5JD4Ca, YH11BCau YH11BCad) was in possession of absolute advantage, at the same time, there were a certain amount of acetic acid and caproic acid. The study found that the lactic acid in calcified pit mud was about 7 times higher than quality pit mud in *Yanghe* base, the *Shuanggou* base was about 11 times. Of course, it could found a strange phenomenon, the calcified pit mud had more pyruvic acid and quality pit mud basically does not had this kind of material. This was very worthy of our thinking.

**Table 1.** Comparison of main organic acid ion contents in different pit muds

| Sample | mg/g |  |  |  |  |  |  |  |  |
| --- | --- | --- | --- | --- | --- | --- | --- | --- | --- |
|  | Lactic acid | Acetic acid | Formic acid | Propionic acid | Isobutyric acid | Butyric acid | Pentanoic acid | Caproic acid | Pyruvic acid |
| S5JD1 | 43.21 | 2.89 | 1.24 | 0.46 | 0.2 | 0.33 | 0.53 | 5.84 | 0.12 |
| S5JB1 | 34.62 | 2.12 | 0.89 | 0.49 | 0.88 | 0.16 | 0.24 | 4.91 | 0.24 |
| S5JD3 | 50.16 | 3.76 | 0.39 | 0.25 | 0.35 | 0.43 | 0.18 | 2.54 | 0.18 |
| S5JD4Ca | 477.28 | 2.14 | 0.35 | 0.35 | 0.52 | 0.23 | 0.28 | 4.68 | 30.35 |
| YH1JN | 25.49 | 3.21 | 0.72 | 0.48 | 0.69 | 0.42 | 0.2 | 10.53 | 0.16 |
| YH11JD | 33.17 | 2.42 | 1.02 | 0.31 | 0.64 | 0.65 | 0.47 | 15.26 | 0.32 |
| YH11BCad | 190.01 | 5.19 | 0.87 | 0.43 | 0.57 | 0.59 | 0.51 | 4.67 | 20.23 |
| YH11BCau | 182.16 | 3.26 | 0.48 | 0.32 | 0.49 | 0.83 | 0.24 | 5.29 | 25.35 |

### The metal ions analysis of the two kinds pit mud

In order to understand more thoroughly of calcification problem, the experiment were tested metal ions of pit mud. Twelve kinds of main organic acids were identified, the specific results were shown in table 2. From the table, calcified pit mud had higher Ca, Mg and Fe metal ions than quality pit mud, *shuanggou* base content were higher than *yanghe* base. Through the calculation, it could found that for the *yangh*e base, the calcium ions, magnesium ions and iron ions of the calcified pit mud was about 17 times, 12 times and 5 times than quality pit mud, respectively. For the *shuanggou* base, it was about 26 times, 23 times and 107 times, respectively. For the other metal ions, there were no significant difference between the two kinds of pit mud.

**Table 2.** Comparison of main metal ions contents in different pit muds

| Sample | mg/g |  |  |  |  |  |  |  |  |  |  |  |
| --- | --- | --- | --- | --- | --- | --- | --- | --- | --- | --- | --- | --- |
|  | Ba | Ca | Mn | Mg | Na | Al | Zn | K | Fe | Cu | P | Pb |
| S5JD1 | 0.42 | 1.49 | 0.35 | 1.03 | 12.04 | 51.72 | 0.04 | 44.7 | 0.73 | 0.04 | 0.19 | 0.02 |
| S5JB1 | 0.45 | 1.67 | 0.33 | 1.06 | 12.11 | 54.18 | 0.03 | 35.67 | 0.78 | 0.03 | 0.3 | 0.02 |
| S5JD3 | 0.28 | 1.59 | 0.5 | 1.98 | 16.03 | 44.11 | 0.04 | 47.96 | 0.6 | 0.04 | 0.19 | 0.02 |
| S5JD4Ca | 0.46 | 39.28 | 0.3 | 24.25 | 11.71 | 40.73 | 0.04 | 36.61 | 75.6 | 0.04 | 0.32 | 0.02 |
| YH1JN | 0.39 | 1.32 | 0.6 | 1.26 | 14.49 | 54.7 | 0.04 | 52.64 | 0.88 | 0.04 | 0.39 | 0.03 |
| YH11JD | 0.23 | 1.92 | 0.42 | 1.5 | 14.53 | 35.67 | 0.05 | 48.13 | 0.94 | 0.05 | 0.29 | 0.02 |
| YH11BCad | 0.42 | 17.43 | 0.57 | 14.98 | 10.78 | 47.96 | 0.04 | 36.41 | 4.19 | 0.05 | 0.22 | 0.02 |
| YH11BCau | 0.42 | 16.54 | 0.48 | 12.22 | 12.78 | 36.61 | 0.06 | 52.03 | 5.58 | 0.04 | 0.41 | 0.02 |

### DNA gel electrophoresis diagram of pit mud

Through the gel electrophoresis diagram, the fragment of prokaryotes was 298 bp and the eukaryotes was 603 bp, it was consistent with the target segments respectively. So it can speculate that the extracted DNA was effective. The specific results were shown in Fig 1. Extracts of microbial genome as templates, with 16 s rDNA V4 area and ITS1-2 area as the target, it could be building high throughput sequencing library. Library stripe size around 350 bp, in line with expectations. The concentration and purity of the library which detected by the Q -bit and Real -time PCR can satisfy the requirements of computer.

**Fig. 1 A:**
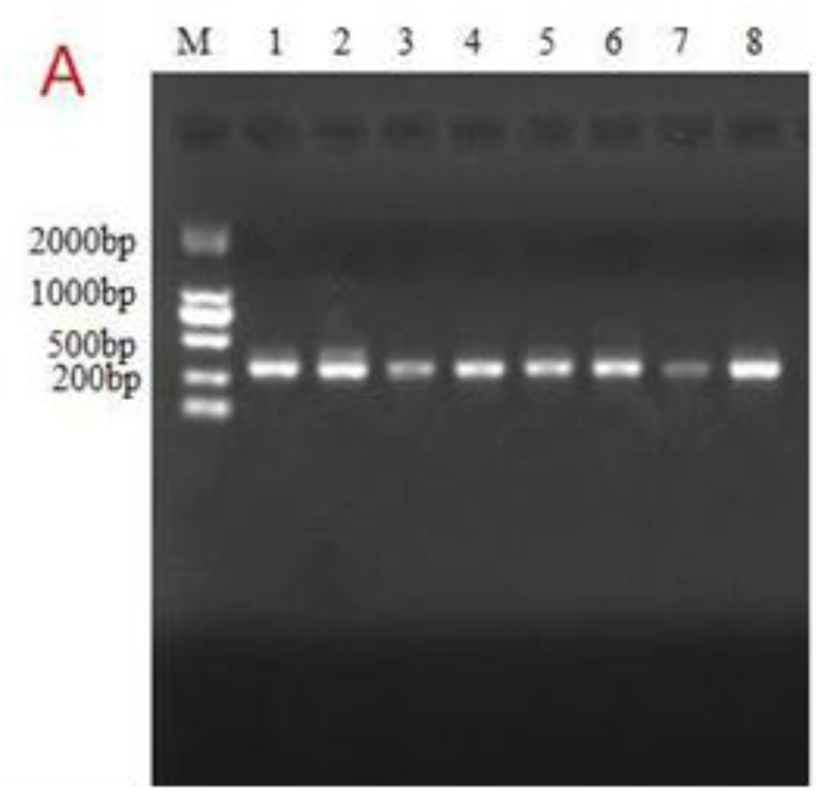
The gel electrophoresis diagram of the prokaryotes.

**Fig. 1 B:**
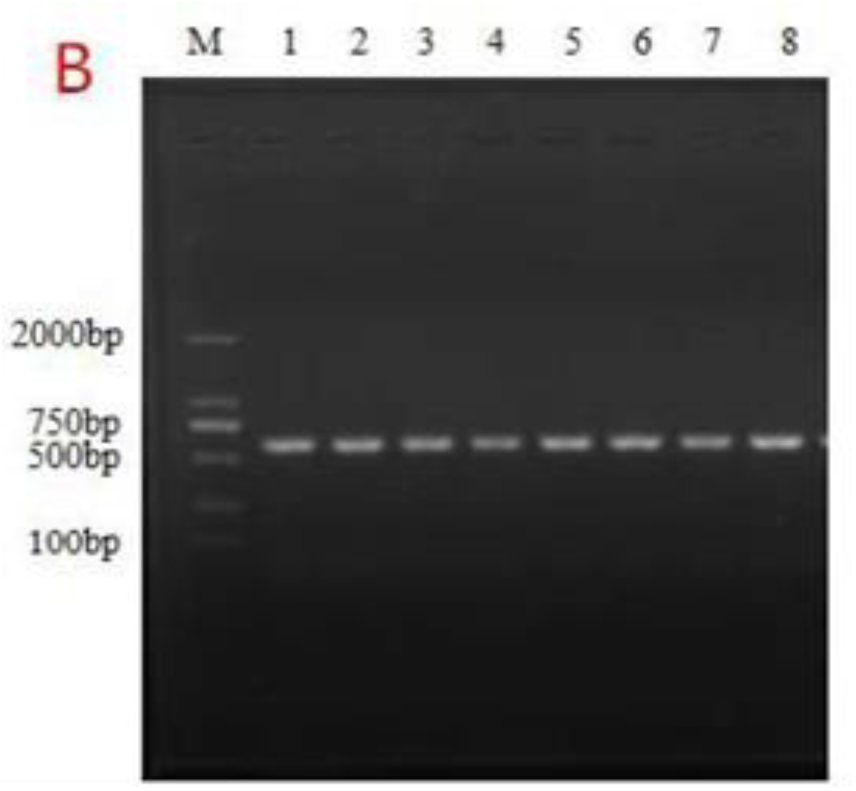
The gel electrophoresis diagram of the eukaryotes

### The OTU result of pit mud and Alpha diversity analysis

Qualified sequencing libraries (including bacteria and fungi) would be for high-throughput sequencing. From the OTU result (Fig 2), it can be found that the Total Tags of calcified pit mud were less than the quality pit mud obviously which both prokaryote microbes and eukaryotic microbes showed the same trend. Those shown that after the calcification of pit mud, lacking of nutrients and too much harmful metabolites, which would be not conducive to microbial habitat. Of course, from the Fig 2, it could be found the difference between the prokaryotes and eukaryotes obviously. The former is almost 8 times than the latter. So it could be guess that the main role of microbes in pit mud would be prokaryotes.

**Fig. 2:**
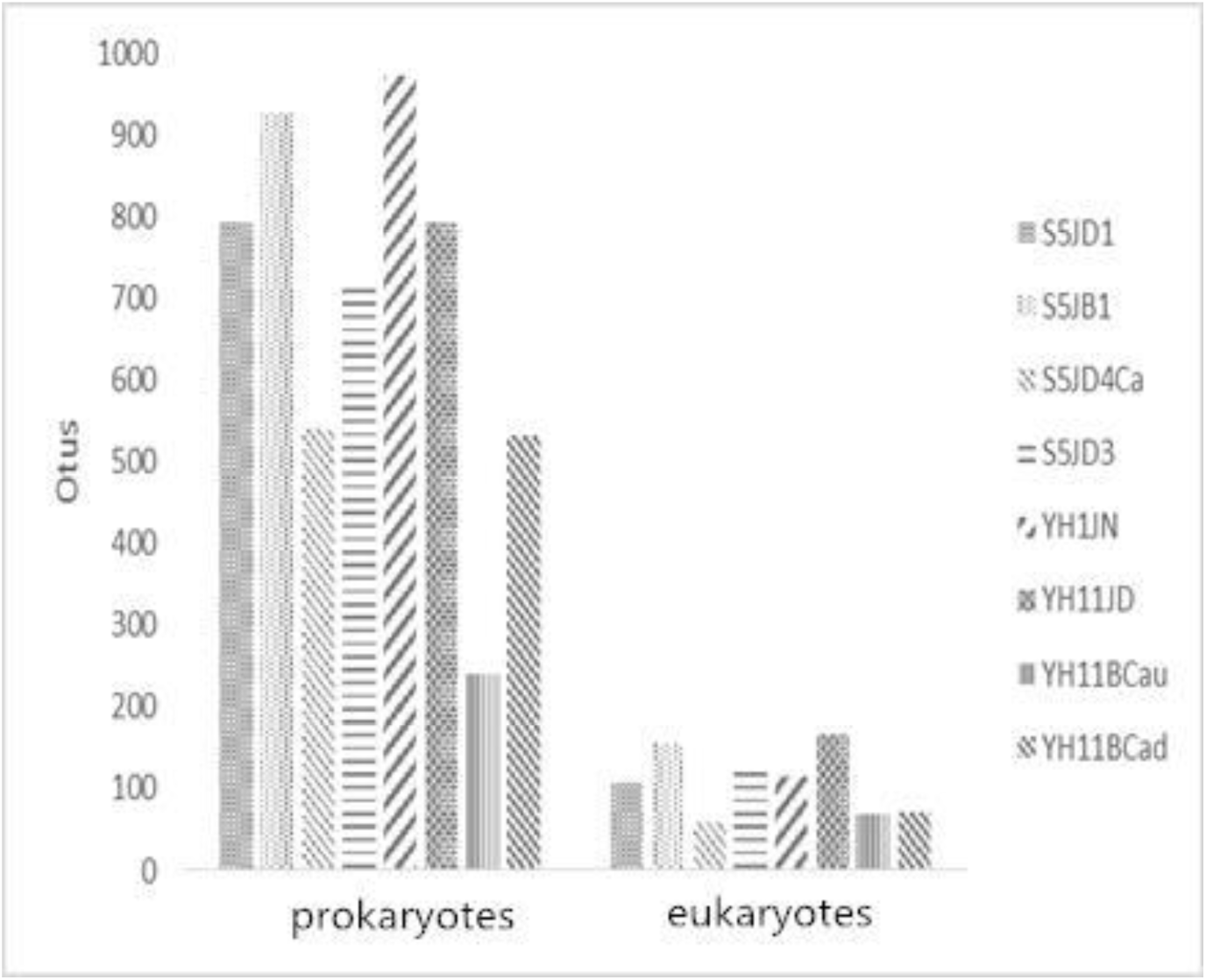
The OTU result of pit mud

In order to detect the reliability and comprehensiveness of data, we made alpha diversity analysis. From the rarefaction curve of Shannon, it can be found that the curves was gradually tends to horizontal line, therefore with the increasing depth of the sample sequence, the number of OTU had reached saturation. Those shown that the depth of sequencing had been basically covered all the microbial species.

### The structure analysis of microbial community

The data analysis shown that the samples were detected prokaryotes 908 strains and the eukaryotes 226 strains. Compared with the quality pit mud, for the *yanghe* manufactory, the prokaryotes reduced 28.2% and the eukaryotes reduced 44.3% in the calcified pit mud. For the *shuanggou* manufactory, the prokaryotes reduced 44.3% and the eukaryotes reduced 80.9%. We hypothesized that the calcified phenomenon of *shuanggou* was more serious than *yanghe*, which was consistent with the facts. From the top sixteen of prokaryotic microorganisms (Fig 3), it could found that the main prokaryotes genera were *Ruminococcaceae, Lactobacillus* and *Prevotella,* but the calcified pit mud had more *Lactobacillus* or *Prevotella,* especially *yanghe* manufactory. For *yanghe* manufactory, the total proportion of *Lactobacillus* and *Prevotella* of quality pit mud and calcified pit mud were 4.5% and 46.5%, respectively. The study found that even in calcification of pit mud, there were still a lot of *Methanocorpusculum,* especially the calcification sample of S5JD4Ca. This was very strange, because traditional expert thought only quality aged pit mud had large amounts of methane bacteria *(17).*

**Fig. 3:**
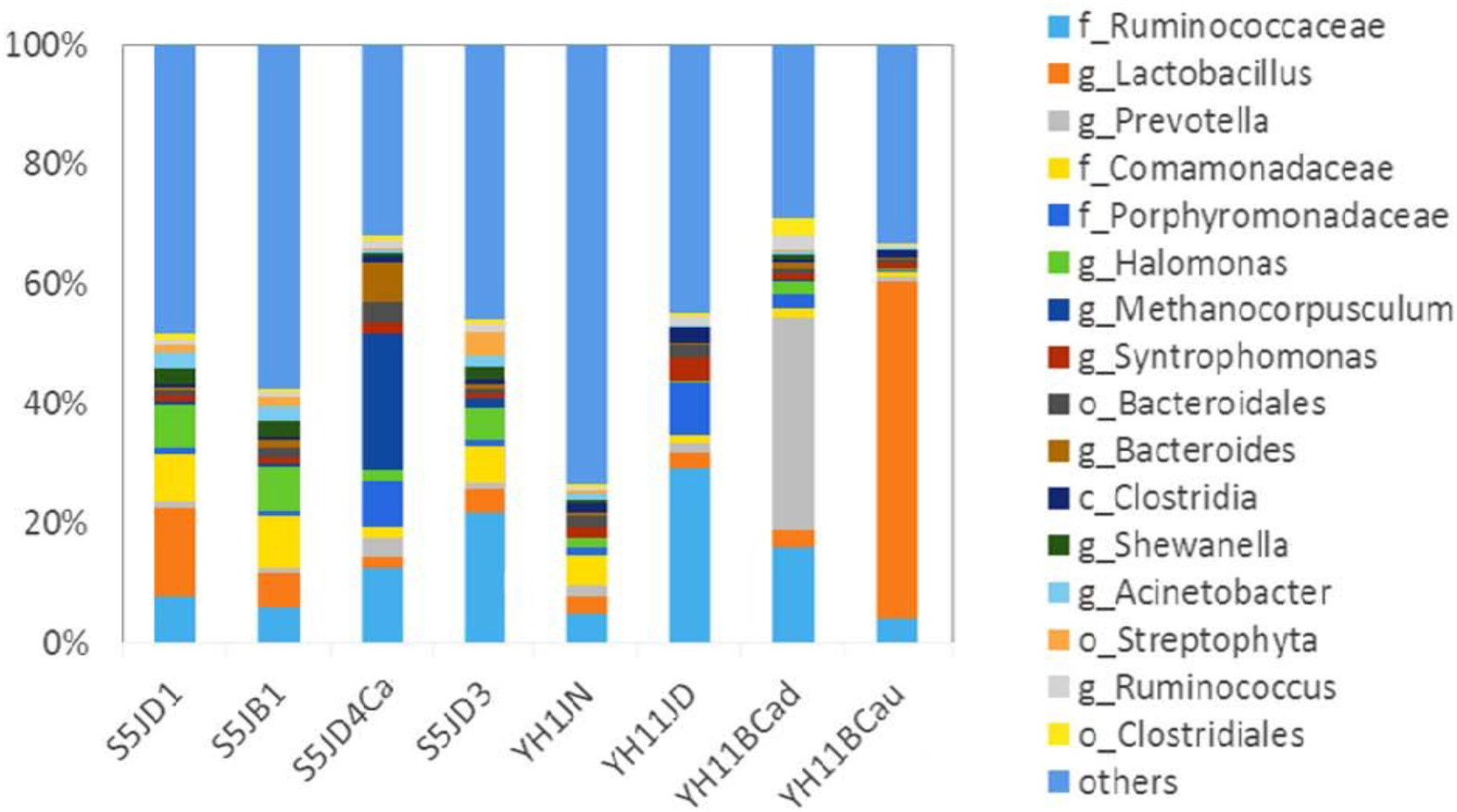
The relative proportions of different prokaryotic microorganisms in different pit muds

From the top ten of eukaryotic microorganisms (Fig 4), it could found that the main eukaryotic genera was *Trichocomaceae,* but the content in calcified pit mud was relatively little. There were also some differences between the *shuanggou* and *yanghe* manufactory, it could found the latter contains more rich microorganisms. For *shuanggou* manufactory, the *Trichocomaceae* proportion of quality pit mud and calcified pit mud were 16.9% and 0.7%, respectively. For *yanghe* manufactory, it were 44.9% and 14.1%, respectively. So it was the objective reflect that the calcification of *shuanggou* pit mud was more serious.

**Fig. 4:**
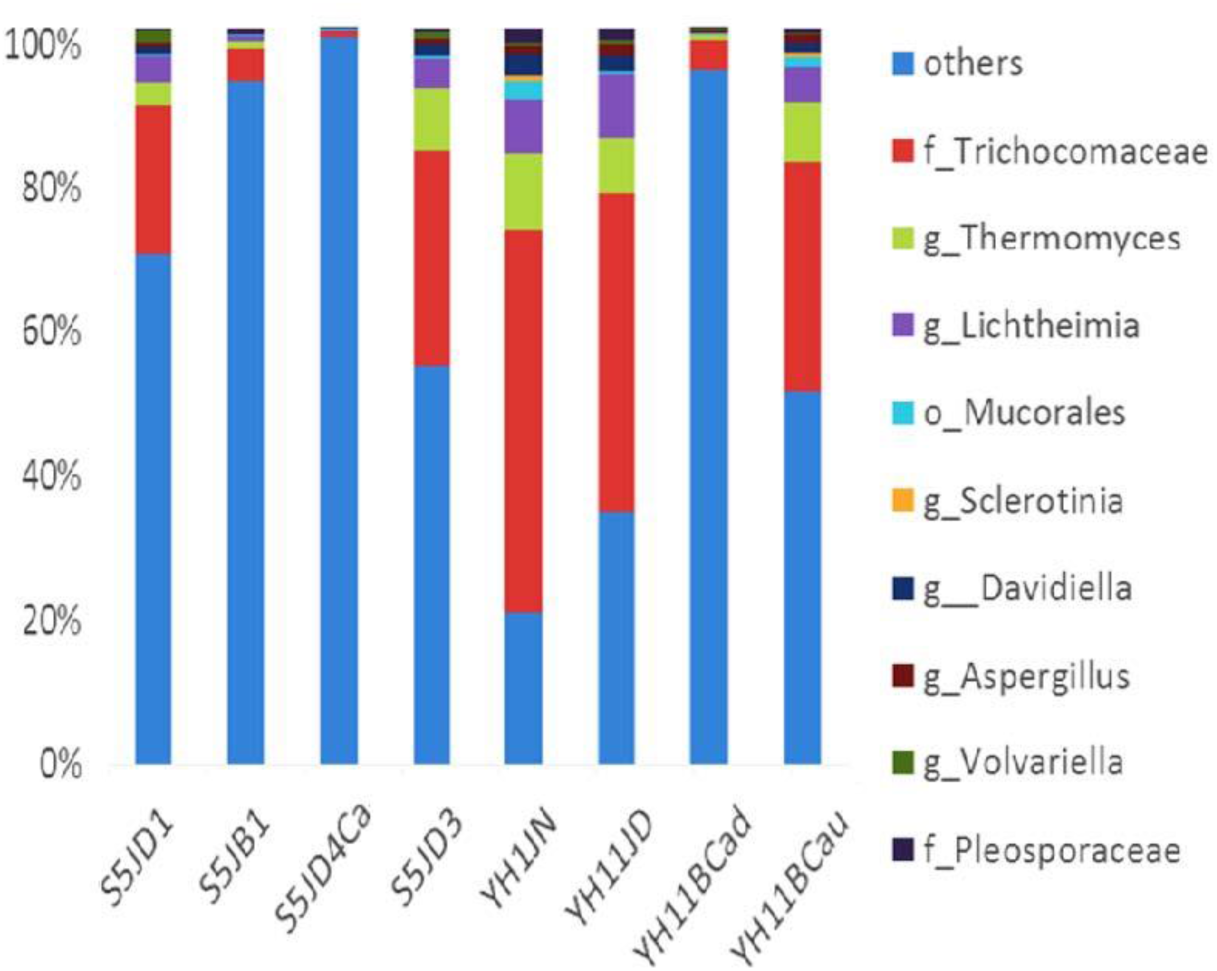
The relative proportions of different eukaryotic microorganisms in different pit muds

## Discussion

Chinese liquor is one of the distilled liquor with global reputation. It is called Chinese culture gem because of its unique traditional fermentation process and a long history of drinking culture. The strong-flavor liquor was occupied about 78% of the Chinese liquor market. Chinese strong liquor plays an important role in Chinese daily life due to it can warm reception guests. The characteristic flavour-aroma formation in Chinese liquor depends on the fermentation style employed, the fermentation foundation (pit mud) and the strains of microorganisms represented *(18).* Therefore, understanding the physical and chemical properties and microbial distribution, characteristics and functionality of pit mud is of importance to understand and found the reason of calcification and improve the quality of Chinese strong liquor. Here, we present the first microbiological analysis in pit mud using high-throughput sequencing and the first time to study the reason of calcification comprehensively.

### The analysis of physical and chemical properties

Through the pit mud contrast between y*anghe* and *shuanggou*, for the organic acids, the results shown that the calcified pit mud had a huge increase in lactic acid content (increased 7 times and 11times, respectively), especially the *shuanggou* manufactory. Its content is far more than the normal brewing system. However, in comparison with the two base pit mud, there had not too big difference. For the metal ions, the results shown that the calcified reason of *yanghe* base pit mud was mainly calcium ions (increased 17 times), the calcified reason of *shuanggou* base pit mud were mainly iron ions and calcium ions (increased 107 times and 26 times, respectively), the content of iron ions was much higher than other pit mud and higher than the content of calcium ion. Of course the magnesium ions was also the main cause of calcification in pit mud.

By analysing the physical and chemical properties (organic acid and metal ions) of the calcified and quality pit mud, it could found that the calcified pit mud contained more Fe^3+^, Mg^2+^, Ca^2 +^ and lactic acid, so it can be guess that those metal ions will combine with lactic acid and form chemical compounds such as calcium lactate, Iron lactate and magnesium lactate. According to the previous studies, the white crystal of calcified pit mud were ferrous lactate, magnesium lactate and copper lactate *(19),* which was also consistent with our study. Those compounds will affects the growth of microbes in pit mud. Here noted an interesting phenomenon, the calcified pit mud had more pyruvic acid and quality pit mud basically does not had this kind of material. The pyruvic acid is a key intermediate in cellular metabolic pathways and it mainly from the metabolic process. However, someone found some defects bacteria can produce pyruvic acid, such as *Acinetobacter, Enterobacter* and *Torulopsis (20).* So the reason of pyruvic acid appeared in calcified pit mud may be the inbalance of the microbial metabolic process.

### The analysis of microbial structure

Because of the complexity of the pit mud, and it found prokaryotes 908 strains and the eukaryotes 226 strains, it was analyzed the main typical microorganism. So we study the three species (*Ruminococcaceae, Lactobacillus* and *Trichocomaceae*). *Ruminococcaceae* was a thermophilic and cellulolytic microorganism. It usually appeared in the intestinal microbiota which was play an important role in the metabolic processes. Through degraded cellulose it can produce such as carbon dioxide, hydrogen, ethanol, acetate. *(21). Lactobacillus* was a genus of Gram-positive facultative anaerobic or microaerophilic rod-shaped bacteria. Many *Lactobacillus* operate using homofermentative metabolism, and some species of *Lactobacillus* use heterofermentative metabolism. The *Lactobacillus* can resistant to acid and need very few nutrients to grow (22). Because of the calcified pit mud had the low pH and the poor nutrition, but the quality pit mud were just the opposite *(23)* and lactic acid bacteria had lower requirements for environmental. So it can be understand that the calcified pit mud had more *Lactobacillus*. As a result it could be guess that the main cause of calcification in pit mud were the several stains, especially *Lactobacillus,* which was consistent with the study of Tao Y *(24)* in pit mud by denaturing gradient gel electrophoresis (DGGE). As for the *Trichocomaceae,* it can inhibit certain bacteria breeding, when these fungi decreased (calcified pit mud was reduced by 24 times), it would lead to excessive breeding of bacteria, even produce large organic acid *(25).*

At the same time, the study found that even in calcification of pit mud, there were still a lot of methane bacteria. But the traditional industry thought only high quality aged pit mud had large amounts of methane bacteria and it was a characteristic metabolites *(17).* So our study may be overthrew the former judging and it is worthy of study late.

### The speculation of the calcification mechanism

The rarefaction curve (in this study, which drown by the Shannon curve) were used to evaluate whether the amount of sequencing was enough to cover all taxa and indirectly reflect the species richness in the sample. The rarefaction curve was measured on the 16S rDNA or 18S rDNA sequence that was the relative proportion of known various OTU and calculated the expectations of OTU number when we extracted the reads of the number of n (n was less than the measured total reads sequence), then according to the value of n and the relative expectation, we drawn the rarefaction curve. From the alpha diversity analysis, these results indicated that our study was meaningful and can be represented the true reflection about microbial diversities and structures in calcified pit mud and quality pit mud. So the result was credible and reliant.

### The speculation of the calcification mechanism

Through the result it can be safely drawn the conclusion that the reason of calcification was complex and regional, for example, the calcification of *yanghe* manufactory was the three kinds of microorganism, especially the *Lactobacillus* and the *Prevotella* (in total increased 11 times), it can produce a large number of organic acid, especially the lactic acid, which combined with the excess metal ions, mainly includes calcium ion, magnesium ion and iron ion (the physical and chemical testing: the calcified pit mud were more than 17, 12 and 5 times higher than the quality pit mud, respectively), the calcification of *shuanggou* manufactory was the imbalance of the microbial community and the structure was changed, which made the microbial metabolic disorders. Those imbalance of microbes lead to produce large amounts of lactic acid in pit mud, which combined with the excess metal ions, mainly includes calcium ion, magnesium ion and iron ion (the physical and chemical testing: the calcified pit mud were more than 26, 23 and 107 times higher than the quality pit mud, respectively).

Taken all together, through the physical and chemical properties and using the method of high-throughput sequencing, we could get a lot of information between the calcified pit mud and the quality pit mud. It can be guess that the microbial structures of calcified pit mud were out of balance, especially the higher *Lactobacillus,* and *Prevotella* and lower *Trichocomaceae,* it would lead to produce a large amount of lactic acid. Those lactic acid will combine with the redundant metal ions, such as calcium ions, iron ion and magnesium ion, and would form the compounds of lactic acid salt. Those compounds would affect the structure of the microbe. It will be a vicious circle. So in order to ease the calcification phenomenon, it can properly reduce the content of those compounds and inhibit the growth of *Lactobacillus* and *Prevotella,* especially the *Lactobacillus*. These may be an effective method. In order to alleviate or solve the problem of calcification, it can be added a food additive in pit mud to, such as hop acid leaching solution and Nisin, those food additive can inhibit the growth of *Lactobacillus* effectively. Due to the calcification had a lower pH value *(26),* it can adjusted by using the method of adding alkali bacteria and balance the microbial community, in the end it can alleviate the calcification of pit mud. Of course that was only my idea, the feasibility of those measures, it still need to carry on the practice and the effect of these substances would also consider the taste of wine.

To the best of our knowledge, none of the previous studies on the microbial comparison between the calcified and quality pit mud. It was the first system analysis for the calcified pit mud and quality pit mud by using high-throughput sequencing. So it provides an exemplary role for revealing the composition of microbial structure in pit mud. This study also provided a good comparison of physical and chemical properties between the calcified pit mud and the quality pit mud, which was significant for the maintenance of pit mud and the quality of liquor. It was also the first time to explore the reason of calcification in pit mud and provides technical and theoretical support for Chinese strong flavor liquor.

## Acknowledgements

This research was supported by National Natural Science Foundation of China (31401680).

